# Genetic ablation of adhesion ligands averts rejection of allogeneic immune cells

**DOI:** 10.1101/2023.10.09.557143

**Authors:** Quirin Hammer, Karlo Perica, Hanna van Ooijen, Rina Mbofung, Pouria Momayyezi, Erika Varady, Karen E. Martin, Yijia Pan, Mark Jelcic, Brian Groff, Ramzey Abujarour, Silje Krokeide, Tom Lee, Alan Williams, Jode P. Goodridge, Bahram Valamehr, Björn Önfelt, Michel Sadelain, Karl-Johan Malmberg

## Abstract

Allogeneic cell therapies hold promise for broad clinical implementation, but face limitations due to potential rejection by the recipient immune system. Silencing of beta-2-microglobulin (*B2M*) expression is commonly employed to evade T cell-mediated rejection, although absence of *B2M* triggers missing-self responses by recipient natural killer (NK) cells. Here, we demonstrate that deletion of the adhesion ligands *CD54* and *CD58* on targets cells robustly dampens NK cell reactivity across all sub-populations. Genetic deletion of *CD54* and *CD58* in *B2M*-deficient allogeneic chimeric antigen receptor (CAR) T and multi-edited induced pluripotent stem cell (iPSC)-derived NK cells reduces their susceptibility to rejection by NK cells *in vitro* and *in vivo* without affecting their anti-tumor effector potential. Thus, these data suggest that genetic ablation of adhesion ligands effectively alleviates rejection of allogeneic immune cells for immunotherapy.

## INTRODUCTION

Allogeneic cellular immunotherapies have the potential to overcome certain limitations of autologous cell therapies due to the possibility for multiplexed gene editing, robust scalability, and on-demand availability for off-the-shelf treatment^1^. However, the immunogenicity of allogeneic cells and the rejection of the infused cell product by the recipient pose significant challenges, which hinder broad clinical implementation^2^.

Genetic deletion of the beta-2-microglobulin (*B2M*) gene successfully abrogates a major arm of CD8^+^ T cell reactivity by abolishing cell surface expression of human leukocyte antigen (HLA) class I^3^. Correspondingly, deletion of HLA class II transactivator (*CIITA*) leads to absence of HLA class II and thereby precludes CD4^+^ T cell responses^4^. Together, these approaches enable substantial avoidance of T cell-mediated rejection^5^. However, the absence of HLA class I triggers missing-self recognition by recipient NK cells^6^, thus necessitating additional immune-modulating strategies to limit recipient NK cell-mediated rejection.

Formation of the immune synapse is an early event during activation of cytotoxic NK and T cells, and essential for target cell killing^7^. The adhesion ligands CD54 (ICAM-1) and CD58 (LFA-3) engage the adhesion receptors LFA-1 and CD2 respectively, and thereby promote the formation of stable synapses^8^. Loss of CD54 and CD58 has been reported to facilitate escape from immune surveillance in virus-infected cells^9–11^ as well as tumors^12–15^. Thus, we hypothesized that reverse-engineering naturally occurring virus and tumor immune escape mechanisms may mitigate the rejection of allogeneic cell products by recipient immune cells.

Here, we demonstrate that simultaneous deletion of *CD54* and *CD58* on HLA class I^low^ target cells limits NK cell activation and killing universally across all sub-populations of NK cells regardless of their inhibitory receptor repertoire. We further show that genetic ablation of *CD54* and *CD58* in *B2M*-deficient allogeneic CAR T cells and iPSC-derived NK cells renders those cells less susceptible to NK cell rejection, while maintaining their anti-tumor effector potential.

## RESULTS

### Silencing of *B2M* expression renders allogeneic immune cells susceptible to missing-self responses by NK cells

Given that genetic silencing of *B2M* expression is employed to mitigate T cell-driven rejection of allogeneic cell products, we investigated the NK cell response profile against primary immune cells with silenced *B2M* expression. To this end, we first treated T cells with *B2M* siRNA, resulting in almost complete loss of HLA class I surface expression (Figure 1A-C). Next, we modelled rejection of allogeneic immune cells by co-culturing siRNA-treated T cells with NK cells from unrelated donors. Compared to cells treated with non-targeting control siRNA, *B2M* siRNA-treated allogeneic T cells showed strongly enhanced susceptibility to killing by NK cells (Figure 1D, E) and elicited robust degranulation of CD56^dim^ NK cells (Figure 1F, G). Simultaneous assessment of multiple sub-populations of CD56^dim^ NK cells revealed that the enhanced reactivity against HLA class I^low^ allogeneic T cell targets was driven by educated NK cells expressing self-HLA-specific inhibitory receptors (KIR2DL1, KIR2LD3, KIR3DL1, and NKG2A in HLA-C1/C2/Bw4 donors; Figure 1H). In contrast, responses by non-educated NK cells not expressing these receptors were minor (Figure 1H), suggesting that missing-self responses prompt allogeneic rejection upon silencing of *B2M*. In line with the principles of NK cell education^16^, the degranulation against HLA class I^low^ T cells increased with the number of inhibitory receptors expressed on the responding NK cell sub-population (Figure 1I).

**Figure 1.**
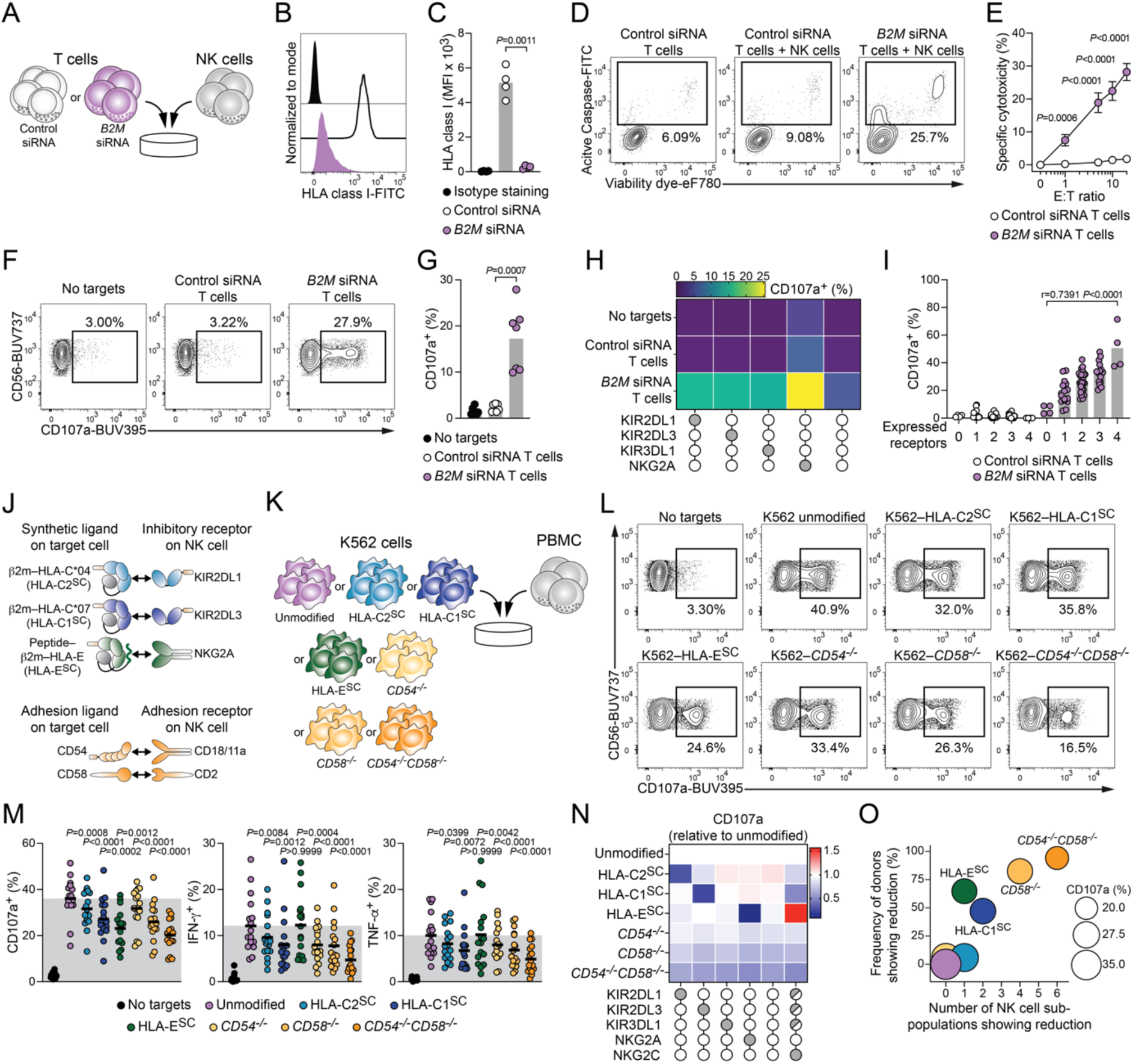
Genetic ablation of *CD54* and *CD58* universally restrains NK cell activity. (A) Schematic illustration of *in vitro* rejection assays between siRNA-treated T cells and NK cells from unrelated healthy donors. (B) Representative HLA class I expression on T cells following siRNA treatment. (C) Summary of HLA class I expression on T cells (n=4 donors in 4 independent experiments). (D) Representative killing of indicated siRNA-treated T cells by NK cells. (E) Summary of specific cytotoxicity (n=7 pairs in 5 independent experiments). (F) Representative degranulation of CD56^dim^ NK cells against indicated siRNA-treated T cells. (G) Summary of CD56^dim^ NK cell degranulation (n=7 donor pairs in 2 independent experiments). (H) Degranulation response pattern of CD56^dim^ NK cell sub-populations stratified for KIR2DL1, KIR2DL3, KIR3DL1, and NKG2A (mean of n=7 donor pairs in 2 independent experiments). (I) Degranulation of CD56^dim^ NK cell sub-populations stratified for number of receptors (n=4 donor pairs in 1 independent experiment). (J) Top: schematic illustration of synthetic HLA molecules as ligands for indicating inhibitory NK cell receptors. Bottom: schematic illustration of adhesion ligands and their cognate receptors. (K) Schematic illustration of *in vitro* NK cell activation experiments between genetically modified K562 target cells and PBMC. (L) Representative degranulation of viable CD14^−^ CD19^−^ CD3^−^ CD56^dim^ NK cells against indicated K562 target cells. (M) Summary of CD56^dim^ NK cell degranulation (left), IFN-γ expression (middle) and TNF-α expression (right). Grey shaded area indicates mean response against unmodified K562 target cells (n=17 donors in 5 independent experiments). (N) Degranulation response of indicated CD56^dim^ NK cell sub-populations against genetically modified K562 target cells relative to unmodified K562 target cells (n=17 donors in 5 independent experiments). (O) Frequency of donors that show reduction plotted against number of CD56^dim^ NK cell sub-populations that show reduction. Bubble size indicates degranulation of CD56^dim^ NK cells. Reduction is defined as relative decrease of at least 25% compared to unmodified K562 cell targets (n=17 donors in 5 independent experiments). C,G: paired t-test. E: Repeated measures two-way ANOVA with Šídák’s multiple comparisons test. I: Pearson correlation. M: Repeated measures one-way ANOVA with Šídák’s multiple comparisons test to unmodified K562 target cells.

To substantiate the data obtained from T cells, we extended our *in vitro* rejection model to allogeneic NK cells serving as targets, where we observed congruent response patterns (Supp. Figure 1A-I).

### Genetic ablation of *CD54* and *CD58* universally restrains NK cell activity

Introduction of ligands for inhibitory receptors such as HLA-E fusion molecules^17^ has been proposed to prevent NK cell responses against *B2M*-deficient allogeneic cell products. To curtail activation of educated NK cells, we constructed single-chain dimers of β2m linked to HLA-C*04:01 (HLA-C2^SC^) and to HLA-C*07:01 (HLA-C1^SC^) as synthetic ligands for the inhibitory receptors KIR2DL1 and KIR2DL3, respectively. We also generated a single-chain trimer of β2m, HLA-E*01:01, and the covalently linked HLA-G_3-11_ peptide VMAPRTLFL^17^ (HLA-E^SC^; Figure 1J). We stably expressed these synthetic ligands in K562 cells, which are inherently HLA class I^low^ and thus serve as prototypical targets for NK cell missing-self responses (Supp. Figure 1J). Functional interrogation of NK cell responses against engineered targets (Figure 1K) confirmed a strong missing-self response upon co-culture with unmodified K562 cells and further revealed that synthetic ligands for inhibitory receptors resulted in a moderate reduction in degranulation as well as interferon gamma (IFN-γ) and tumor necrosis factor alpha (TNF-α) expression in CD56^dim^ NK cells (Figure 1L, M).

Since cell adhesion and immune synapse formation are critical early events in NK cell activation, we next directly compared introduction of synthetic ligands for inhibitory receptors to genetic ablation of the key adhesion ligands *CD54* and *CD58* (Figure 1 J). Here, we found that absence of *CD54* or *CD58* on the target cells limited NK cell responses and dual ablation of both *CD54* and *CD58* resulted in marked reduction of all tested NK cell effector functions that was more pronounced than the effects of synthetic ligands for inhibitory receptors (Figure 1L, M). Closer inspection of NK cell response patterns at the sub-population level exposed that, as expected, expression of synthetic ligands mediated partial reduction due to specific inhibition of sub-populations expressing the respective cognate inhibitory receptor (Figure 1N). Furthermore, introduction of HLA-E^SC^ potently inhibited NKG2A-expressing NK cells, but simultaneously elicited robust effector function of NK cells expressing the HLA-E-specific activating receptor NKG2C, thereby nullifying overall inhibition (Supp. Figure 1J, K). In contrast, deletion of *CD54* and *CD58* affected NK cell activity across all tested sub-populations, independent of inhibitory receptor expression and including the NKG2C^+^ sub-population (Figure 1N), likely because NK cells unanimously require adhesion for immune synapse formation and activation. We next assessed the number of NK cell sub-populations that showed reduced activity against a given K562 target and determined the degree to which the different edits affected overall NK cell responses. This analysis further confirmed a pervasive effect of dual *CD54* and *CD58* deletion in all 6 tested NK cell sub-populations and uncovered reduced overall NK cell responses in 16 out of 17 donors (Figure 1O). Together, these data demonstrate that genetic ablation of the adhesion ligands *CD54* and *CD58* universally restrains NK cell responses against HLA class I^low^ targets.

### *CD54*^−/−^*CD58*^−/−^ target cells are resistant to NK cell attack due to curtailed adhesion

Since ablation of *CD54* and *CD58* resulted in broadly muted NK cell functions, we explored whether this effect was indeed mediated by decreased adhesion to target cells and whether it results in diminished killing of target cells. To this end, we performed conjugate formation assays between K562 targets and NK cells (Figure 2A). Compared to unmodified K562 cells, a significantly lower frequency of NK cells formed conjugates with *CD54^−/−^CD58^−/−^*targets (Figure 2B, C). Accordingly, lower conjugate formation translated to significantly less pronounced killing of *CD54^−/−^CD58^−/−^* targets compared to their unmodified counterparts (Figure 2D, E). We extended these assays with single cell analyses using micro-well chips that enable tracking of individual NK cells over time (Figure 2F, G)^18,19^ and seeded chips with unmodified or *CD54^−/−^CD58^−/−^* K562 targets, followed by addition of low numbers of NK cells to obtain wells containing a single NK cell. When following individual NK cells from the same donor, we found that they spent less time in contact with *CD54^−/−^CD58^−/−^* targets than with unmodified K562 cells (Figure 2H, Supp. Video 1). Additionally, NK cells that engaged *CD54^−/−^CD58^−/−^* targets made less committed contacts (Supp Figure 2A, B). The reduced contact was similarly reflected in a lower proportion of single NK cells killing adhesion ligand-deficient targets (Figure 2I).

**Figure 2.**
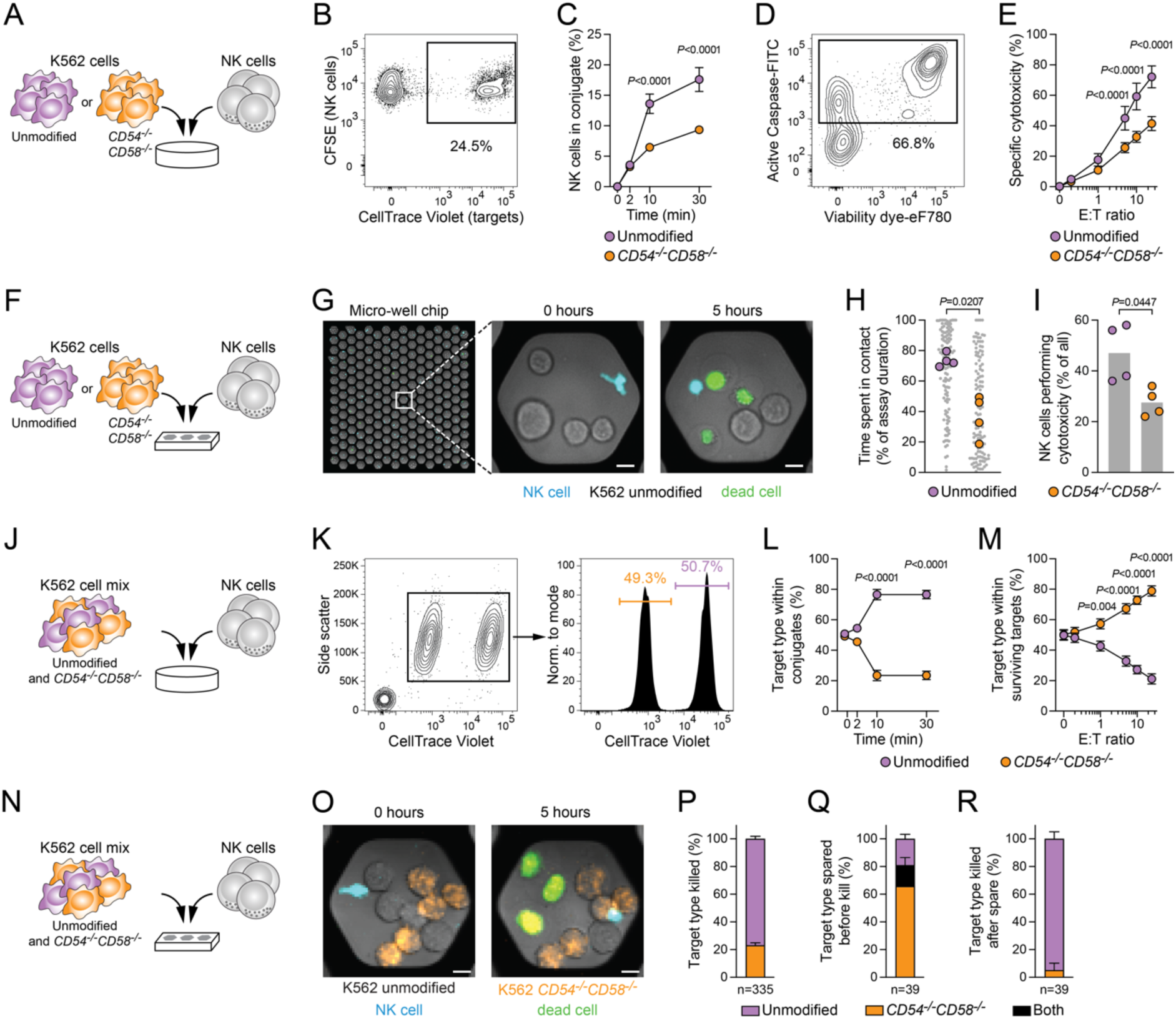
*CD54*^−/−^ *CD58*^−/−^ target cells are resistant to NK cell attack due to curtailed adhesion. (A) Schematic illustration of conjugation and cytotoxicity assays between K562 cells and NK cells. (B) Representative conjugate formation of CFSE-labelled NK cells to CellTrace Violet-labelled target cells. (C) Summary of conjugate formation (n=8 donors in 3 independent experiments). (D) Representative killing of target cells by NK cells. (E) Summary of specific cytotoxicity (n=8 donors in 4 independent experiments). (F) Schematic illustration of *in vitro* microwell assays between K562 cells and NK cells. (G) Representative display of co-cultures between single NK cells and multiple K562 cells over time. Scale bar 10 μm. (H) Summary of relative time of NK cells spent in contact with target cells (n=4 donors in 3 independent experiments). (I) Summary of relative NK cell cytotoxicity (n=4 donors in 3 independent experiments). (J) Schematic illustration of *in vitro* conjugation and cytotoxicity assays between mixed K562 cells and NK cells. (K) Representative deconvolution of mixed target cells based on CellTrace Violet intensity. (L) Summary of distribution of target cell type within formed conjugates (n=8 donors in 5 independent experiments). (M) Summary of distribution of target cell type within surviving target cells in cytotoxicity assays (n=8 donors in 4 independent experiments). (N) Schematic illustration of *in vitro* microwell assays between mixed K562 cells and NK cells. (O) Representative display of co-cultures between single NK cells and multiple mixed target cells over time. Scale bar 10 μm. (P) Summary of distribution of target cell type killed in first cytotoxic event (n=335 individual NK cells; n=4 donors in 3 independent experiments). (Q, R) Sequential killing of mixed target cells by single NK cells. (Q) Distribution of target cell types with which single NK cells form contacts but do not kill prior to killing another target cell (n=39 sparing events prior to a kill; n=4 donors in 3 independent experiments). (R) Distribution of target cell types which single NK cells kill after having spared another target cell (n=39 killing events following a spare; n=4 donors in 3 independent experiments). C, E, L, M: Repeated measures two-way ANOVA with Šídák’s multiple comparisons test. H, I: Paired t-test.

We examined NK cell responses in a competition assay by labelling target cells with different concentrations of CellTrace Violet and mixing them at a 1:1 ratio, followed by addition of NK cells (Figure 2J, K). Analyses of the distribution of target cell types in conjugate assays revealed that the vast majority of NK cells formed conjugates with unmodified K562 cells, while *CD54^−/−^CD58^−/−^* targets were detected only in a minority of conjugates (Figure 2L). Likewise, in competition cytotoxicity assays, unmodified K562 cells were preferentially killed by NK cells and thereby significantly depleted from the surviving target cell population, whereas viable *CD54^−/−^CD58^−/−^* targets were enriched (Figure 2M). Moreover, when exposed to a mix of unmodified and *CD54*^−/−^ *CD58*^−/−^ target cells in micro-well chips (Figure 2N, O), individual NK cells mediated their first cytotoxic event predominantly against unmodified target cells and to a much lesser extent against *CD54^−/−^CD58^−/−^*targets (Figure 2P). Finally, sequential analyses of cytotoxic activity against multiple targets showed that individual NK cells favored sparing *CD54^−/−^CD58^−/−^*targets (Figure 2Q) before moving on to lyse primarily unmodified K562 cells (Figure 2R).

These findings highlight that genetic deletion of *CD54* and *CD58* in target cells curtails NK cell adhesion and renders target cells resistant to NK cell-mediated lysis.

### Genetic ablation of *CD54* and *CD58* confers enhanced survival to *B2M^−/−^*allogeneic CAR T cells *in vitro* and *in vivo*

To explore the implications of genetic ablation of *CD54* and *CD58* in a setting of allogeneic cell therapy, we generated T cells expressing a second generation CD19-CAR combined with deletion of *TRAC* to avoid graft-versus-host reactivity^20^. We detected CD54 and CD58 to be expressed on resting CAR T cells and found that expression of both proteins was increased following activation (Figure 3A). We genetically deleted *B2M* alone or *B2M* together with *CD54* and *CD58* in CAR T cells (Figure 3B). Next, we assessed the anti-tumor functionality of engineered CAR T cells against CD19^+^ Nalm-6 targets *in vitro* (Figure 3C) and found that CAR T cells killed tumor cells with comparable efficiency across several effector:target ratios in short-term assays (Figure 3D) and with similar kinetics in long-term assays (Figure 3E). To model the impact of adhesion ligand deletion on NK cell rejection *in vitro*, we co-cultured a mix of unmodified, *B2M^−/−^*, and *B2M^−/−^CD54^−/−^CD58^−/−^*CAR T cells with NK cells (Figure 3F). Akin to our findings from siRNA-treated T cell targets, deletion of *B2M* resulted in diminished survival compared to unmodified allogeneic CAR T cells (Figure 3G). Importantly, *B2M^−/−^CD54^−/−^CD58^−/−^*allogeneic CAR T cells showed recovered survival in the presence of NK cells relative to their *B2M^−/−^* counterparts (Figure 3G).

**Figure 3.**
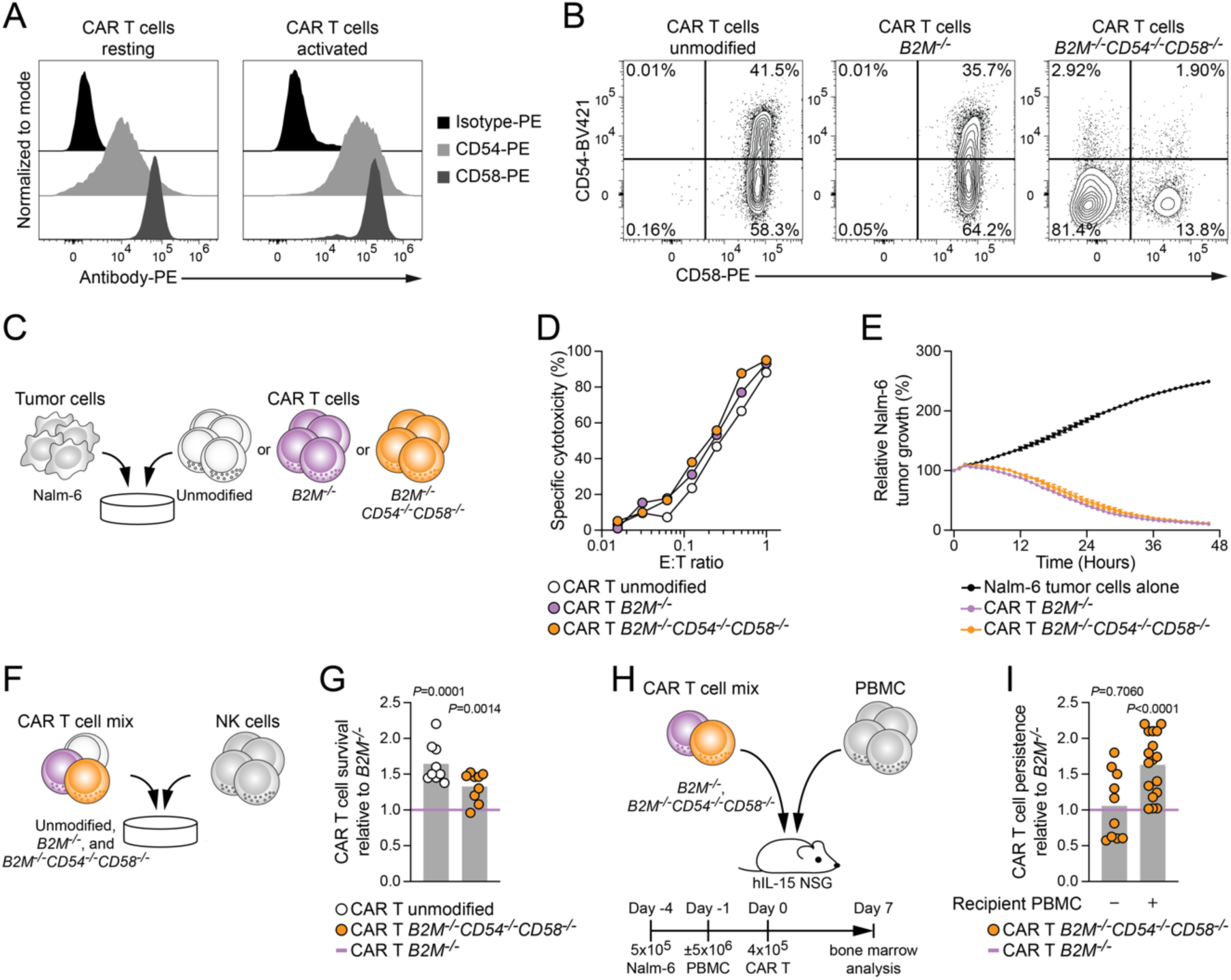
Genetic ablation of *CD54* and *CD58* confers enhanced survival to *B2M*^−/−^ allogeneic CAR T cells in the presence of immune effector cells *in vitro* and *in vivo*. (A) Representative staining of CD54 and CD58 on resting (left) and activated (right) CAR T cells. (B) Representative staining of CD54 and CD58 on activated CAR T cells either unmodified (left), *B2M*-deficient (middle), or combined deletions of *B2M*, *CD54*, and *CD58* (right). (C) Schematic illustration of *in vitro* anti-tumor functional assay of genetically modified CAR T cells. (D) Cytotoxicity of indicated CAR T cells against Nalm-6 target cells for 18 h at varying E:T ratios. (E) Cytotoxicity of indicated CAR T cells against Nalm-6 target cells for 46 h at E:T= 1 (n=1 experiment). (F) Schematic illustration of *in vitro* rejection assays between CAR T cells and NK cells. (G) Summary of CAR T cell survival relative to *B2M*^−/−^ CAR T cells upon co-culture with NK cells (n=9 donors in 3 independent experiments). (H) Schematic illustration of *in vivo* rejection assays between CAR T cells and PBMC engrafted into human IL-15-transgenic NSG mice. (I) Summary of CAR T cell persistence relative to *B2M*^−/−^ CAR T cells 7 days post transfer (n=10 animals without PBMC engraftment and n=17 animals with PBMC engraftment). G, I: One-sample t-test to CAR T *B2M^−/−^*.

We further explored CAR T cell persistence in an *in vivo* model of allogeneic cell rejection by inoculating human IL-15 transgenic immunodeficient mice (hIL-15 NSG) with Nalm-6 tumor cells, followed by engraftment of recipient PBMC and transfer of a mix of *B2M^−/−^* and B*2M^−/−^CD54^−/−^CD58^−/−^* allogeneic CAR T cells (Figure 3H). Seven days after transfer, *B2M^−/−^CD54^−/−^CD58^−/−^*CAR T cells displayed significantly enhanced persistence relative to *B2M^−/−^*CAR T cells with intact adhesion ligands in animals engrafted with recipient PBMC (Figure 3I). In line with our observation from K562 cells as targets, *B2M*^−/−^ CAR T cells engineered to express HLA-E^SC^ were partially protected from attack by NK cells (Supp. Figure 3A), albeit this was not efficient against NK cells harboring sizable NKG2C^+^ sub-populations (Supp. Figure 3B, C).

### Multi-edited *CD54*^−/−^*CD58*^−/−^ allogeneic iPSC-derived NK cells display resistance to NK cell rejection *in vitro* and *in vivo*

Multiplex editing in primary T cells remains challenging^21^ and raises concerns for genomic instability^22^. These shortcomings can be attenuated by employing iPSC as a next-generation platform for allogeneic therapy^23^. Thus, we engineered iPSC with multiple modalities including a CD19-CAR, a IL-15/IL-15 receptor alpha fusion protein (IL-15/IL-15Rα), and a high-affinity non-cleavable variant of CD16 (hnCD16)^24–27^, subsequently referred to as unmodified. Subsequently, we extended the multi-editing approach by genetic deletion of either *B2M* and *CIITA* alone or *B2M* and *CIITA* in conjunction with deletion of *CD54* and *CD58*. Employing a clonal master cell bank as a source of engineered iPSC for down-stream differentiation and expansion of iPSC-derived NK cells resulted in end products displaying highly homogenous phenotypes (Figure 4A, B; Supp. Figure 4A-D,). Comparing the anti-tumor functionality of unmodified, *B2M^−/−^CIITA^−/−^*, and *B2M^−/−^CIITA^−/−^CD54^−/−^CD58^−/−^*iPSC-derived NK cells against K562 target cells demonstrated near identical degranulation and expression of IFN-γ as well as TNF-α (Figure 4C-F). These functional similarities were additionally reflected in comparable polyfunctional responses of iPSC-derived NK cells (Supp. Figure 4E). Moreover, we observed almost indistinguishable expression of IFN-γ in response to stimulation with the innate cytokines IL-12, IL-15, and IL-18, independent of whether the NK cells were differentiated from unmodified, *B2M^−/−^CIITA^−/−^*, or *B2M^−/−^ CIITA^−/−^CD54^−/−^CD58^−/−^* iPSC (Supp. Figure 4F).

**Figure 4.**
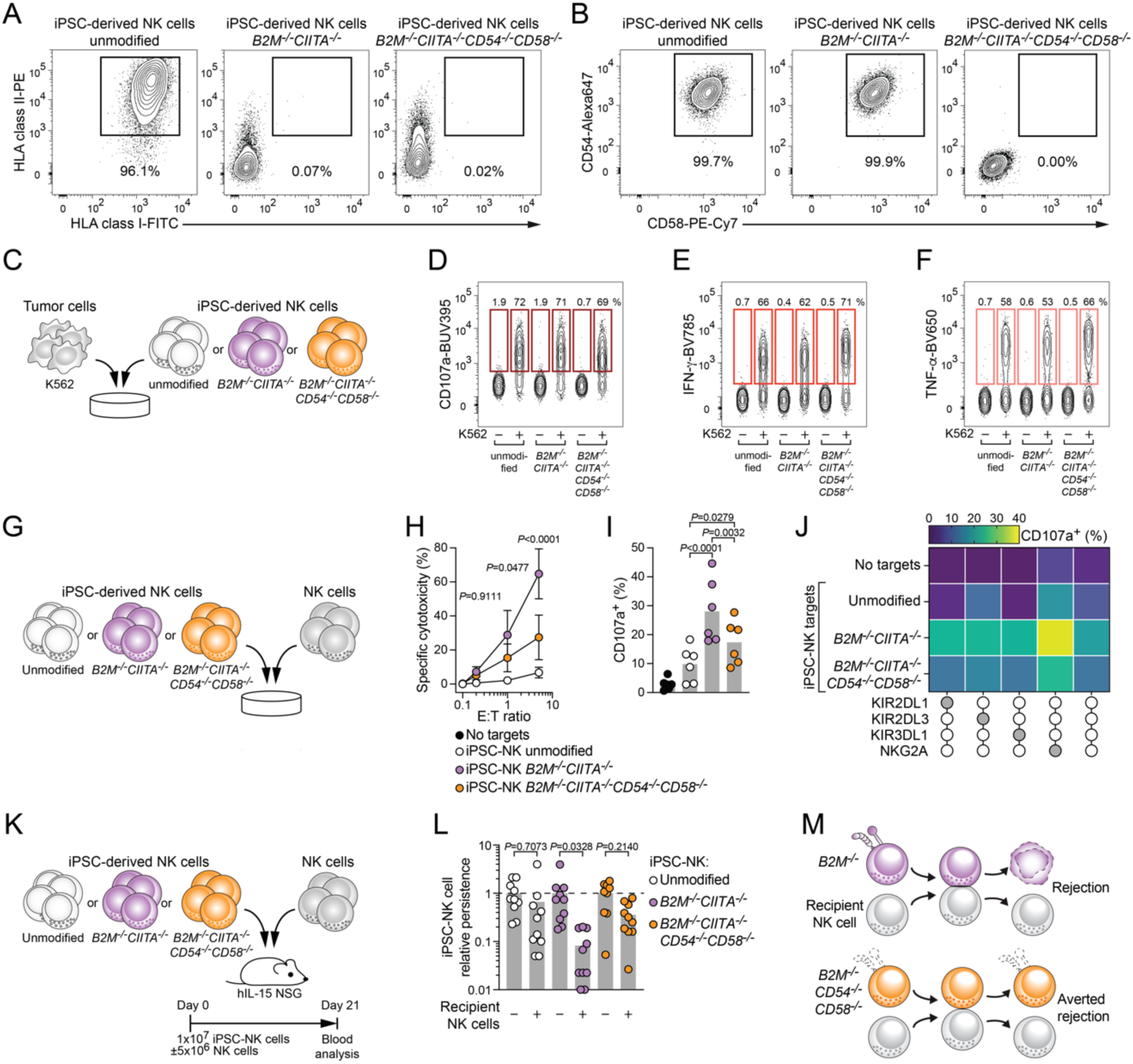
Multi-edited *CD54*^−/−^*CD58*^−/−^ allogeneic iPSC-derived NK cells display resistance to NK cell rejection *in vitro* and *in vivo*. (A) Representative staining of HLA class I and HLA class II on viable CD45^+^ CD56^+^ iPSC-derived NK cells either expressing CD19-CAR, IL-15/IL-15Ra, and hnCD16 alone (unmodified; left), or combined with genetic deletion of *B2M* and *CIITA* (middle), or combined with genetic deletion of *B2M*, *CIITA*, *CD54*, and *CD58* (right). (B) Representative staining of CD54 and CD58 on viable CD45^+^ CD56^+^ iPSC-derived NK cells with indicated genotype. (C) Schematic illustration of *in vitro* anti-tumor functional assay of iPSC-derived NK cells. Representative (D) degranulation, (E) IFN-γ expression, (F) TNF-α expression, of viable CD45^+^ CD56^+^ iPSC-derived NK cells with indicated genotype upon co-culture with K562 tumor target cells (n=1 experiment). (G) Schematic illustration *of in vitro* rejection assays between iPSC-derived NK cells and NK cells. (H) Summary of specific cytotoxicity against the indicated iPSC-derived NK cells (n=4 donors in 2 independent experiments). (I) Summary of CD56^dim^ NK cell degranulation against the indicated iPSC-derived NK cells (n=6 donors in 2 independent experiments). (J) Degranulation response pattern of CD56^dim^ NK cell sub-populations stratified for KIR2DL1, KIR2DL3, KIR3DL1, and NKG2A (mean of n=6 donor pairs in 2 independent experiments). (K) Schematic illustration of *in vivo* rejection assays between iPSC-derived NK cells and activated NK cells engrafted into human IL-15-transgenic NSG mice. (O) Summary of iPSC-derived NK cell persistence relative to animals without recipient NK cell engraftment (n=9-10 animals per group in 2 independent experiments). (P) Schematic illustration of rejection of *B2M*^−/−^ allogeneic cells by recipient NK cells (top) and averting of rejection by combining *B2M*^−/−^ with deletion of CD54 and CD58. H, I: Repeated measures two-way ANOVA with Šídák’s multiple comparisons test. L: Ordinary one-way ANOVA with Šídák’s multiple comparisons test.

We next tested the resistance of iPSC-derived NK cell products to effector NK cells in *in vitro* rejection assays (Figure 4G). Analogous to our data from siRNA-treated cells and CAR T cells, genetic deletion of *B2M* and *CIITA* rendered iPSC-derived NK cells susceptible to attack by NK cells from unrelated donors and, importantly, multi-edited *B2M^−/−^CIITA^−/−^CD54^−/−^CD58^−/−^* iPSC-derived NK cells were significantly less susceptible to killing by primary NK cells (Figure 4H). Correspondingly, degranulation of primary effector NK cells was significantly reduced upon co-culture with *B2M^−/−^CIITA^−/−^CD54^−/−^ CD58^−/−^* iPSC-derived NK cells as compared to *B2M^−/−^CIITA^−/−^*iPSC-derived NK cells serving as targets (Figure 4I), and markedly lower missing-self responses were evident for all educated effector NK cell sub-populations (Figure 4J). When surveilling the persistence of iPSC-derived NK cells over time in co-cultures with PBMC, we found that genetic ablation of *CD54* and *CD58* resulted in consistently sustained survival (Supp. Figure 4G, H). Finally, we examined the persistence of iPSC-derived NK cells in an *in vivo* model of NK cell-mediated rejection. For this, we reconstituted hIL-15 NSG mice with primary NK cells acting as recipient compartment and transferred either unmodified, *B2M^−/−^CIITA^−/−^*, or *B2M^−/−^CIITA^−/−^CD54^−/−^CD58^−/−^*allogeneic iPSC-derived NK cells into the animals (Figure 4K). Three weeks after transfer, *B2M^−/−^CIITA^−/−^* iPSC-derived NK cells were significantly depleted in animals bearing recipient NK cells, whereas multi-edited *B2M^−/−^CIITA^−/−^CD54^−/−^CD58^−/−^*iPSC-derived NK cells displayed considerable resistance to rejection (Figure 4L).

Collectively, these findings indicate that genetic ablation of the adhesion ligands *CD54* and *CD58* effectively averts rejection of *B2M^−/−^CIITA^−/−^*allogeneic immune cells (Figure 4M).

## DISCUSSION

Rejection by the recipient is considered a central limitation of allogeneic cell therapies. In this study, we demonstrate that deletion of *CD54* and *CD58* in HLA class I^low^ allogeneic immune cells effectively reduces rejection by NK cells *in vitro* and *in vivo*. These findings support the concept that unidirectional manipulation of the immune synapse can be leveraged to generate hypoimmunogenic cells with extended persistence.

Allogeneic cells adoptively transferred for cancer immunotherapy require sufficient persistence to mediate their anti-tumor functions, similar to autologous cells products which show a correlation between persistence and clinical efficacy^28^. Rejection by the recipient’s immune system restrains the persistence of allogeneic cells and thereby limits their clinical value, especially regarding future multi-dosing regimens and off-the-shelf availability. Thus, extending the persistence has the capacity to markedly enlarge the therapeutic window of allogeneic immunotherapy.

The primary trigger for rejection upon transfer across HLA barriers are the HLA molecules themselves. Decreased expression of HLA class I and II is required to avoid T cell-mediated rejection and is achieved by deletion of *B2M* and *CIITA*^29,30^. However, the absence of HLA class I render cells vulnerable to missing-self recognition by recipient NK cells^6^. We established a reductionist model of allogeneic rejection that uses siRNA treatment to diminish *B2M* expression^31^ and observed that resting NK cells responded robustly to HLA class I^low^ primary T and NK cells. Educated NK cells expressing NKG2A or self-HLA class I-specific KIR displayed predominant effector activity and their responsiveness increased with the number of receptors expressed. Thus, NK cell-mediated rejection of allogeneic HLA class I^low^ cells is driven by a *bona fide* missing-self response^6^ and adheres to the principles of NK cell education^16^.

Based on these observations, it seems reasonable to limit NK cell rejection by introducing ligands for educating receptors, such as synthetic variants of HLA-E^17,32^ or HLA-C, into allogeneic cells prior to adoptive transfer. The use of HLA-like synthetic proteins presents a fundamental challenge due to the existence of paired inhibitory and activating NK cell receptors recognizing the identical ligand. One such pair are the inhibitory NKG2A and the activating NKG2C receptors, which both bind to HLA-E/peptide complexes^33^. Notably, NKG2C is a signature marker of adaptive NK cells, which lack NKG2A and expand in response to cytomegalovirus (CMV) infection^34–36^. Depending on CMV seroprevalence, up to 40% of all individuals habor sizeable populations of NKG2A^−^ NKG2C^+^ adaptive NK cells^35^, and transfer of allogeneic cells expressing synthetic HLA-E ligands into these individuals is likely to be detrimental due to NKG2C-driven enhancement of rejection. In line with previous data^17^, we found that single chain trimers of β2m, HLA-E, and a peptide epitope efficiently inhibited NKG2A-expressing NK cells. However, the net effect at the overall CD56^dim^ level was nullified due to a simultaneous stimulatory effect on NKG2A^−^ NKG2C^+^ NK cells. Thus, approaches relying on HLA-like molecules are highly dependent on the relative size of the inhibited NK cell sub-population and at the same time carry the risk of potentially activating another sub-population of NK cells in the recipient, hampering broad implementation.

The adhesion ligands CD54 and CD58 contribute to the earliest events during synapse formation between NK cells and their targets^7,8^. Loss-of-function of CD54 and CD58 is detrimental to NK cell activity in diverse settings. As such, HIV-infected cells suppress CD54 surface expression, thereby escaping from NK cell killing^10^. Moreover, CMV-infected cells, which down-regulate HLA class I to avoid CD8^+^ T cell recognition, evade NK cell immunosurveillance by retaining CD58 intracellularly^11^. Paralleling viral escape, tumors such as B cell lymphomas may down-regulate both CD54 and CD58^12^. Mutations in *CD58* and copy number loss thereof are negative prognostic markers in lymphoma^13,14^ and the majority of tumors lacking HLA class I display a concurrent absence of CD58^15^, indicative of selection in response to missing-self recognition by NK cells. We therefore hypothesized that manipulating adhesion ligands would generate resistance to NK cells despite absence of HLA class I. Our data demonstrate that deletion of *CD54* and *CD58* in allogeneic *B2M^−/−^CIITA^−/−^* CAR T and iPSC-derived NK cells improved survival in presence of NK cells. Importantly, this approach was independent of NK cell sub-populations and operational in the vast majority of NK cell donors, suggesting potential for widespread implementation.

Alternative strategies to limit NK cell-mediated rejection by over-expressing CD47^37^ or equipping transferred cells with alloimmune defense receptors^38^ have been reported to demonstrate preclinical efficacy. These strategies take advantage of the inhibitory receptor SIRP1α on activated NK cells^39^ and exploit the direct elimination of activated, 4-1BB-expressing NK cells^38^, respectively, and represent approaches that intervene at later stages of the NK cell activation cascade, subsequent to immune synapse formation.

Specifically tailoring the immune synapse to achieve a desired functional outcome has recently been applied to amplify anti-tumor responses of CAR NK cells^40^. In this setting, introduction of a scaffolding domain augmented synapse formation between CAR NK cells and tumor cells, resulting in notably improved tumor control by fine-tuning adhesion properties^40^.

Overall, modulation of the immune synapse emerges as a promising strategy to optimize the functional properties and persistence of immune cells for adoptive transfer. Our findings highlight that naturally occurring escape strategies can serve as blueprints for augmenting cell therapy and support the concept that genetic ablation of adhesion ligands such as CD54 and CD58 enables manipulation of the immune synapse to avert rejection of allogeneic immune cells.

### Limitations of the study

This study reports the deletion of adhesion ligands as conceptual advance to reduce rejection by NK cells. We focused on mitigating NK cell activity since lack of HLA class I promotes NK cell responses. It is well established that allograft rejection involves multiple vital players including T cells, B cells, NK cells, and macrophages but also antibodies and the complement system. Therefore, continuous efforts are required to achieve complete hypoimmunogenicity that comprehensively down-modulates other components of rejection such as induction of donor-specific antibodies or complement activity. Moreover, extended profiling of the anti-tumor functionality mediated by adhesion ligand-deficient effector cells is a prerequisite for further application.

## ACKNOWLEDGEMENTS

This project has received funding from the European Union’s Horizon 2020 research and innovation programme under the Marie Sklodowska-Curie grant agreement No. 8382909 (to QH). This work was supported by the Swedish Research Council (223310), the Swedish Children’s Cancer Society (PR2020-1059), the Swedish Cancer Society (21-1793Pj), Sweden’s Innovation Agency, and the Karolinska Institutet. The work was further supported by the Research Council of Norway (275469, 237579), Center of Excellence: Precision Immunotherapy Alliance (332727), the Norwegian Cancer Society (190386, 223310), The South-Eastern Norway Regional Health Authority (2021-073), EU H2020-MSCA Research and Innovation programme (801133), Knut and Alice Wallenberg Foundation, Swedish Foundation for Strategic Research, the US National Cancer Institute (P01 CA111412, P009500901, all to KJM).

## AUTHOR CONTRIBUTIONS

Conceptualization: QH, KP, JPG, BV, MS, KJM.

Methodology: QH, KP, HvO, RM, PM.

Formal analysis: QH, KP, HvO.

Investigation: QH, KP, HvO, RM, PM, EV, RA, YP, KEM, MJ, BG, AW.

Resources: SK, TL.

Writing – original draft: QH, KJM.

Writing – Review & Editing: All authors.

Supervision: QH, BÖ, MS, KJM.

Funding acquisition: QH, MS, KJM.

## DECLARATION OF INTERESTS

KJM is a consultant and has research support from Fate Therapeutics. KJM has research support from Oncopeptides. KJM and QH are consultants at Vycellix. All relationships has been approved by Oslo University Hospital, University of Oslo and Karolinska Institute. RM, EV, YP, MJ, BG, RA, TL, AW, JPG and BV are employees at Fate Therapeutics. MSK has licensed IP (unrelated to this study) to Fate Therapeutics and MS receives research support (for unrelated studies) from Fate Therapeutics.

## SUPPLEMENTARY FIGURE LEGENDS

**Supplementary Figure 1. Silencing of *B2M* expression renders allogeneic NK cells susceptible to missing self responses by NK cells from unrelated donors**. (A) Schematic illustration of *in vitro* rejection assays between siRNA-treated NK cells and NK cells. (B) Representative HLA class I expression on NK cells following siRNA treatment. (C) Summary of HLA class I expression on NK cells (n=8 donors in 4 independent experiments). (D) Representative killing of indicated siRNA-treated NK cells by NK cells. (E) Summary of specific cytotoxicity (n=8 pairs in 6 independent experiments). (F) Representative degranulation of CD56^dim^ NK cells against indicated siRNA-treated NK cells. (G) Summary of CD56^dim^ NK cell degranulation (n=9 donor pairs in 2 independent experiments). (H) Degranulation response pattern of CD56^dim^ NK cell sub-populations stratified for KIR2DL1, KIR2DL3, KIR3DL1, and NKG2A (mean of n=9 donor pairs in 2 independent experiments). (I) Degranulation of CD56^dim^ NK cell sub-populations stratified for number of receptors (n=4 donor pairs in 1 independent experiment). (J) Representative expression of synthetic ligands for inhibitory receptors on K562 cells (left) and expression of the adhesions ligands CD54 and CD58 (right). (K) Representative degranulation of viable CD14^−^ CD19^−^ CD3^−^ CD56^dim^ NKG2A^+^ NKG2C^−^ (left) and NKG2A^−^ NKG2C^+^ (right) NK cells against indicated K562 target cells. (L) Correlations between NKG2C^+^/NKG2A^+^ ratio with CD56^dim^ NK cells and functional responses (CD107a^+^, IFN-γ^+^, and TNF-α^+^) of either K562-HLA-E^SC^ (green) or K562-*CD54^−/−^CD58^−/−^* (orange) targets relative to unmodified K562 cells (n=17 donors in 5 independent experiments).

C,G: paired t-test. E: Repeated measures two-way ANOVA with Šídák’s multiple comparisons test. I, L: Pearson correlation.

**Supplementary Figure 2. NK cells fail to engage in committed contacts with *CD54*^−/−^ *CD58*^−/−^ target cells.** (A) Representative display of a non-committed (left) and a committed (right) contact of NK cells with K562 targets. Scale bar 5 μm. (B) Summary of committed contacts against the indicated K562 targets (n=5 donors in 4 independent experiments).

B: Paired t-test.

**Supplementary Figure 3. HLA-E^SC^ renders *B2M^−/−^*CAR T cells susceptible to NKG2C^+^ NK cells**. (A) Summary of CAR T cell survival relative to *B2M*^−/−^ CAR T cells upon *in vitro* co-culture with NK cells containing less than 10% NKG2C^+^ cells (n=6 donors in 2 independent experiments). (B) Summary of CAR T cell survival relative to *B2M*^−/−^ CAR T cells upon *in vitro* co-culture with NK cells containing more than 10% NKG2C^+^ cells (n=3-4 donors in 3 independent experiments). (C) Summary of CAR T cell persistence relative to *B2M*^−/−^ CAR T cells upon transfer into human IL-15-transgenic NSG mice (n=8-10 animals per group).

A-C: One sample t-test to *B2M^−/−^* CAR T cells.

**Supplementary Figure 4. Multi-edited *CD54*^−/−^*CD58*^−/−^ allogeneic iPSC-derived NK cells retain their functionality and display resistance to rejection *in vitro***. (A) Schematic illustration of the iPSC-to-NK cell differentiation and expansion process. (B) Representative expression of pluripotency markers SSEA-4 and TRA-81-1 on viable iPSC with indicated genotypes. (C) Representative expression of the NK cell markers CD45 and CD56 on viable CD14^−^ iPSC-derived NK cells following differentiation and prior to expansion *in vitro*. (D) Fold expansion of iPSC-derived NK cells with the indicated genotypes during co-culture with K562–4-1BBL–mbIL-21 feeder cells *in vitro* for 14 days (n=1-2 expansions). (E) Polyfunctionality of indicated iPSC-derived NK cells upon co-culture with K562 tumor target cells (n=1 experiment). (F) IFN-γ expression of the indicated iPSC-derived NK cells following exposure to IL-12, IL-15, and IL-18 for 24 h (n=1 experiment). (G) Schematic illustration of *in vitro* rejection assays between iPSC-derived NK cells and PBMC. (H) Persistence of indicated iPSC-derived NK cells upon co-culture with PBMC relative to iPSC-derived NK cells cultured without PBMC (n=3 donors in 2 independent experiments)

H: Repeated measures two-way ANOVA with Šídák’s multiple comparisons test.

**Supplementary Video 1. Contact and killing dynamics of single NK cells challenged with either *CD54*^−/−^ *CD58*^−/−^ or unmodified K562 cells**

## MATERIALS AND METHODS

### Cells and cell lines

Buffy coats from healthy donors were obtained from the Department of Clinical Immunology and Transfusion Medicine, Karolinska Institute as approved by the Ethical Review Board Stockholm (DNR 2020-05289) or at MSK from purchased from the New York Blood Center (institutional review board-exempted). Cryopreserved PBMC were thawed and NK cells and T cells were purified by magnetic separation using human NK cell Isolation Kit and CD3 MicroBeads, respectively (both Miltenyi Biotec). KIR-ligands of healthy donors were determined using the Olerup SSP KIR HLA Ligand kit (CareDx).

K562 and Nalm-6 cells were obtained from ATCC. Cells lines were regularly tested for presence of mycoplasma using either MycoAlert Mycoplasma Detection Kit (Lonza) or sequencing-based Mycoplasmacheck (Eurofins).

All cells were maintained in complete RPMI (RPMI-1640 supplemented with 2 mM glutamine, 10% [V/V] FBS, 100 U/mL penicillin, and 100 μg/mL streptomycin; all Gibco/ThermoFisher) at 37°C and 5% CO_2_.

### Flow cytometry

Flow cytometric analyses were performed following established guidelines^41^. In brief, cell suspensions were incubated with combinations of fluorochrome-conjugated antibodies (Supp. Table 1) at optimized concentrations in PBS for 20 min at room temperature (RT). Viable cells were identified with LIVE/DEAD Fixable Aqua Dead Cell Stain Kit or eBioscience Fixable Viability Dye eFluor 780 (both ThermoFisher). Streptavidin-BV711 (BD Biosciences) was used as secondary reagent in combination with biotinylated antibodies. Surface-stained cells were fixed with BD Cytofix/Cytoperm (BD Biosciences) and permeabilized using BD Perm/Wash Buffer (BD Biosciences) according to the manufacturer’s instructions prior to staining of intracellular cytokines for 30 min 4 °C. Samples were stored in PBS containing 2% (V/V) FBS and 2 mM EDTA until acquisition on an LSR Fortessa flow cytometer (BD Biosciences). Analyses were performed with FlowJo v10.7.1 (BD Biosciences).

### *B2M* silencing in primary immune cells

Expression of *B2M* was silenced using siRNA as previously described^31,42^. In brief, purified T cells or NK cells from donors with C1/C2 Bw4 KIR-ligand status were cultured at 1×10^6^ cells/mL in Accell siRNA medium (Dharmacon) supplemented with 50 U/mL IL-2 (Miltenyi) or 1 ng/mL IL-15 (RnD), respectively, and treated with 1 uM Accell control non-targeting siRNA or *B2M* #13 siRNA (both Dharmacon) for 96 h. After siRNA treatment, HLA class I protein expression was assessed with flow cytometry and the cells were used as targets for purified NK cells from unrelated healthy donors.

### Co-culture of NK cells with siRNA-treated allogeneic immune cells

NK cell responses against allogeneic immune were evaluated using purified NK cells from donors with C1/C2/Bw4^+^ KIR-ligand status as effectors and purified NK or T cells from unrelated matched donors with the corresponding KIR-ligand status as targets. For cytotoxicity assays, siRNA-treated target NK or T cells were labelled with 2 uM CellTrace Violet (Thermo Fisher), treated with 1 uM Concanamycin A (Santa Cruz) for 30 minutes, washed extensively with complete RPMI, and subsequently co-cultured with effector NK cells at varying effector:target ratios for 6 hours. Caspase activity was determined with Cleaved Caspase-3 Staining Kit (Abcam) and viability of target cells was assessed using eBioscience Fixable Viability Dye eFluor 780 (ThermoFisher). Target NK cells were identified as single CellTrace Violet^+^ CD3^−^ CD56^dim^ lymphocytes. Specific cytotoxicity was calculated as follows: (% dead target cells in experimental condition – % dead target cells in spontaneous control) / (100% – % dead target cells in spontaneous control) × 100.

For degranulation assays, 1×10^5^ CellTrace Violet-labelled siRNA-treated target NK or T cells were co-cultured with 1×10^5^ effector NK cells in the presence of anti-CD107a antibody for 6 hours. GolgiStop and GolgiPlug (both BD) were added after one hour to inhibit cytokine secretion. After five more hours of co-culture, cell suspensions were stained for flow cytometric analysis. Effector NK cells were identified as single viable CellTrace Violet^−^ CD3^−^ CD56^dim^ lymphocytes and further stratified as indicated.

### Genetic engineering of K562

Synthetic HLA-like ligands were designed based on previous reports to engage HLA-specific NK cell receptors^17^. The following fusion proteins were designed: HLA-C1 single-chain dimer (HLA-C1^SC^) consisting of *B2M* and *HLA-C07:01* fused by a (G_4_S)_4_ linker, HLA-C2 single-chain dimer (HLA-C2^SC^) consisting of *B2M* and *HLA-C04:01* fused by a (G_4_S)_4_ linker, and HLA-E single-chain trimer (HLA-E^SC^) consisting of the HLA-G_3-11_ peptide VMAPRTLFL, *B2M*, and *HLA-E*01:01* fused by (G_4_S)_3_ and (G_4_S)_4_ linkers, respectively.

Fusion protein expression cassettes were cloned into the lentiviral expression vector LeGO-G2 (kind gift from Boris Fehse, Addgene plasmid #251917; http://n2t.net/addgene:25917; RRID: Addgene_25917). Lentiviral particles were generated by transfection of Lenti-X 293T cells (Takara) with 15 ug LeGO expression vector, 10 ug pRSV-Rev (kind gift from Didier Trono (Addgene plasmid #12253; http://n2t.net/addgene:12253; RRID:Addgene_12253), 15 ug pMDLg/pRRE (kind gift from Didier Trono (Addgene plasmid #12251; http://n2t.net/addgene:12251; RRID:Addgene_12251), and 5 ug pCMV-VSV-G (kind gift from Bob Weinberg (Addgene plasmid #8454; http://n2t.net/addgene:8454; RRID:Addgene_8454) using Lipofectamine 3000 (Invitrogen). Lentiviral particles were harvested, concentrated with Lenti-X Concentrator (Takara), used to transduce K562 cells with an MOI of 20. Genetically engineered K562 cells were sorted for high transgene expression using a MA900 cell sorter (Sony). HLA-C1^SC^ and HLA-C2^SC^ were detected using recombinant human KIR2DL3-Fc and KIR2DL1-Fc fusion proteins (both RnD), respectively, along with goat anti-human IgG Fc secondary antibody (polyclonal, ThermoFisher). HLA-E^SC^ was detected with an anti-HLA-E antibody (clone 3D12, BioLegend). For genetic deletion of CD54 and/or CD58, gRNA (IDT, Supp. Table 2) were mixed with Cas9 protein (IDT), followed by RNP electroporation using the Nucleofector II device (Lonza). Genetically modified K562 cells were maintained in complete RPMI and routinely monitored for transgene expression.

### NK cell functional assays

To test NK cell responses against genetically modified K562 target cells, 1×10^6^ PBMC were co-cultured with 1×10^5^ K562 cells in V-bottom 96-well plates in the presence of anti-CD107a and anti-NKG2C antibodies (Supp. Table 1). GolgiStop and GolgiPlug (both BD) were added after one hour to inhibit cytokine secretion. After five more hours of co-culture, cell suspensions were stained for flow cytometric analysis including fixation, permeabilization, and intracellular staining for cytokines. NK cells were identified as single viable CD14^−^ CD19^−^ CD3^−^ CD56^dim^ lymphocytes.

To determine sub-population response patterns, degranulation was assessed for each NK cell sub-population relative to the sub-population’s degranulation against unmodified K562 cells.

To analyze the frequency of donors showing a reduction and the number of NK cell sub-populations showing a reduction, a reduction in degranulation was defined as at least 25% lower degranulation against a given genetically modified K562 target as compared to unmodified K562 cells.

### Conjugation assays of NK cells against genetically modified K562 cells

Unmodified and *CD54^−/−^CD58^−/−^* K562 cells were labeled with 2 uM CellTrace Violet (ThermoFisher) and purified NK cells were labeled with 2 uM CFSE (ThermoFisher). 5×10^5^ target cells and 2×10^5^ NK cells were combined in round-bottom tubes, centrifuged at 20 *g* for 2 min, and subsequently incubated at 37°C for the indicated time periods. The cells were fixed immediately with pre-warmed Cytofix/Cytoperm solution (BD) and stored on ice. Conjugates were analyzed by determining the frequency of CellTrace Violet^+^ events within CFSE^+^ NK cells. Competition conjugation assays were performed by labeling unmodified K562 cells and *CD54*^−/−^*CD58*^−/−^ K562 cells with 2 uM and 0.2 uM CellTrace Violet (ThermoFisher), respectively, and analyzing the frequencies of the differentially labeled cells within CFSE^+^ CellTrace Violet^+^ conjugates.

### Cytotoxicity assays of NK cells against genetically modified K562 cells

Unmodified K562 cells and *CD54^−/−^CD58^−/−^* K562 cells were labeled with 2 uM CellTrace Violet (ThermoFisher) and co-cultured with purified NK cells at varying effector:target ratios. Caspase activity was determined with Cleaved Caspase-3 Staining Kit (Abcam) and viability of target cells was assessed using eBioscience Fixable Viability Dye eFluor 780 (ThermoFisher). Specific cytotoxicity was calculated as follows: (% dead target cells in experimental condition – % dead target cells in spontaneous control) / (100% – % dead target cells in spontaneous control) × 100. Competition cytotoxicity assays were performed by labeling unmodified K562 cells and *CD54*^−/−^*CD58*^−/−^ K562 cells with 2 uM and 0.2 uM CellTrace Violet (ThermoFisher), respectively, and analyzing the frequencies of the differentially labeled cells within the surviving population of viability dye^−^ active Caspase^−^ CellTrace Violet^+^ target cells.

### Micro-well experiments and imaging

Micro-well experiments were conducted using previously described silicon-glass micro-well chips^43^. Two types of chips were used: hexagonal 60 µm-wide wells for live, single NK cell tracking experiments, and square 350 µm-wide wells for confocal analysis of contact quality. Purified NK cells were cultured in complete RPMI supplemented with 100 U/mL IL-2 (Peprotech) overnight and labelled with CellTrace Yellow (ThermoFisher) according to the manufacturer’s instructions before being seeded into the micro-well chip.

For single NK cell contact dynamics and killing analysis, labelled NK cells were seeded with unstained unmodified or *CD54*^−/−^*CD58*^−/−^ K562 cells in microwells. Co-cultures were imaged every 3 min for 15-16 hours using a widefield microscope (Zeiss Axio Observer). Only wells with a single NK cell and a minimum of 3 target cells were included in the analysis, and wells in which the NK cell did not engage any target throughout the assay were excluded. NK cells were considered engaged in target cell contact if the two cells overlapped for at least two consecutive frames, and target cell killing events were detected using SYTOX Green (ThermoFisher) and Incucyte Caspase-3/7 Dye (Sartorius).

For single NK cell competition killing analysis, labelled NK cells were seeded with a mix of unmodified and *CD54*^−/−^*CD58*^−/−^ K562 cells, where either the unmodified or the *CD54*^−/−^*CD58*^−/−^ cells were labelled with CellTrace FarRed (ThermoFisher). Co-cultures were imaged every 15 min for 15 hours using a confocal microscope (Zeiss LSM 880). Only wells with a single NK cell and a minimum of 2 unmodified as well as 2 *CD54*^−/−^ *CD58*^−/−^ cells were included in the analysis. Wells with a difference larger than 1 cell between unmodified and *CD54*^−/−^*CD58*^−/−^ K562 cells were excluded to avoid bias in selective killing. Data was pooled from different staining combinations to account for staining effects.

To analyze contact quality, labelled NK cells and unstained target cells (unmodified or *CD54*^−/−^*CD58*^−/−^ K562 cells) were seeded in microwells and allowed to interact for 80 minutes. The cells were processed in-well and fixed using Cytofix Fixation buffer, permeabilized with Perm/Wash buffer (both BD), blocked with 5% (V/V) FBS in PBS, as well as stained with anti-α-Tubulin antibody (Sigma Aldrich) and with phalloidin– Abberior STAR 635 (Sigma Aldrich) to visualize F-actin. Imaging was performed using a confocal microscope (Zeiss LSM 880), and image analysis was conducted using

CellProfiler (Broad Institute) and ImageJ (NIH). The synapse region between NK cells and target cells was determined by analyzing the overlapping signal between the two cells and calculating the ratio of synapse size to total NK cell size. NK cell-target cell conjugates were automatically classified as committed with a synapse region of ≥ 20% oft he NK cell perimeter, whereas small ratios were considered as non-committed.

### Retroviral vectors for generation of CAR T cells

Plasmids encoding the SFG γ-retroviral vector^44^ were prepared as previously described^45,46^. VSV-G pseudotyped retroviral supernatants derived from transduced gpg29 fibroblasts (H29) were used to construct stable retroviral-producing cell lines as previously described^47^. Viral supernatants were concentrated 10-fold using RetroX concentrator (Takara).

### Generation and engineering of CAR T cells

Purified T cells were activated with Dynabeads Human T-Activator CD3/CD28 (ThermoFisher) at a 1:1 bead:cell ratio in complete RPMI supplemented with 5 ng/mL IL-7 and IL-15 (both PeproTech) and after 48-72 h, beads were removed. Cas9 protein (Berkeley) was incubated with gRNA (Synthego; Supp. Table 2) of interest at a 1:1 molar ratio for 10 min at 37°C. For multiplex editing, the quantity of each guide used was reduced by 25%. 2×10^6^ or 10×10^6^ cells were resuspended with P3 buffer (Lonza) and mixed with 60 or 300 pmol *TRAC* RNP in a total volume of 20 or 100 μl, respectively. T cells were subsequently transfected by electroporation of RNP using setting E0115 of the Nucleofector II device (Lonza). Cells were cultured overnight at density of 3×10^6^ cells/mL in complete RPMI supplemented with 5 ng/mL IL-7 and IL-15 to account for cell loss after electroporation (∼33%). The next day, T cells were transduced by centrifugation on Retronectin (Takara)-coated plates.

### *In vitro* killing of Nalm-6 tumor cells by CAR T cells

Nalm-6 tumor cells expressing firefly luciferase have been previously described^45,48^ and were used as targets to assess the cytotoxicity of CAR T cells in a luciferase-based assay. Effector and target cells were co-cultured in triplicates at the indicated E:T ratios in complete RPMI for 18 h. Luciferase Bright-Glo substrate (Promega) was added and emitted light was measured using a luminescence plate reader or IVIS Imaging System (Xenogen) and quantified using Living Image software (Xenogen). Cytotoxicity was determined as (1 − (RLUsample)/(RLUmax)) × 100.

### Co-culture of NK cells with allogeneic CAR T cells

NK cell responses against allogeneic CAR T cells were evaluated using purified NK cells from healthy donors as effectors and engineered CAR T cells from unrelated donors as targets. NK cells were labelled with CFSE (ThermoFisher) and 1×10^5^ NK cells were cultured with 5×10^4^ CAR T cells for 8-12 hours. SYTOX™ Blue Dead Cell Stain and CountBright Absolute Counting Beads (both ThermoFisher) were used to enumerate surviving CFSE^−^ CAR T cells. Percent survival was calculated as follows: (CARs in alloreactive well)/(CARs in control well) × (beads in control well)/(beads in alloreactive well) × 100.

### *In vivo* persistence assay of CAR T cells in mice engrafted with recipient immune cells

We used NOD.Cg-Prkdcscid Il2rgtm1Wjl Tg(IL15)1Sz/SzJ (referred to as hIL-15-NSG) mice, which support the engraftment of primary NK cells. For determining the persistence of CAR T cells, 8-to 12-week-old female hIL15-NSG mice were inoculated with 5×10^5^ Nalm-6 cells expressing firefly luciferase-GFP on day –4 by tail vein injection. On day –1, 5×10^6^ thawed PBMC were engrafted by tail vein injection to mimic a recipient immune system and on day 0, 4×10^5^ genetically engineered CAR T cells bearing a mixture of the modifications of interest were infused by tail vein injection. Mice were euthanized on day 7 and bone marrow was collected. Single cell suspension were prepared followed by erythrocyte lysis with ACK buffer (ThermoFisher) and CAR T cells were identified as human CD45^+^ CAR^+^ cells. The relative persistence of CAR T cells in each animal was calculated by comparing the frequency of cells bearing the modifications of interest at day 7 to pre-infusion frequencies.

### Generation, engineering, differentiation, and expansion of iPSC and iPSC-derived NK cells

Human iPSC were generated following previously described protocols^49,50^ and differentiated into hematopoietic progenitors and CD34^+^ cells using established methods^51,52^. In brief, iPSC-derived CD34^+^ cells were transferred to NK cell differentiation B0 medium (2:1 mixture of Dulbecco modified Eagle medium and Ham F12 medium supplemented with 2 mM L-glutamine, 100 U/mL penicillin, and 100 μg/mL streptomycin, 25 uM β-mercaptoethanol, 10% [V/V] human AB serum, 5 ng/ml sodium selenite, 50 uM ethanolamine, 20 mg/ml ascorbic acid [all ThermoFisher], 5 ng/mL IL-3 [first week only; ThermoFisher], 30 ng/mL stem cell factor [SCF ThermoFisher], 20 ng/mL IL-15 [RnD], and 10 ng/mL Fml-like tyrosine kinase 3 ligand [FLT3L; ThermoFisher]). The cells were maintained in these differentiation conditions for 20 days with weekly media changes until they reached the desired CD45^+^ CD56^+^ phenotype. Thereafter, iPSC-derived NK cells were expanded by co-culture with irradiated K562–41BBL–mbIL-21 feeder cells in B0 media supplemented with 250 U/ml IL-2 (Miltenyi) for 14 days. Import of and experiments with iPSC and iPSC-derived cells were approved by the Ethical Review Board Stockholm (DNR 2020-00750).

### Functional assays of iPSC-derived NK cells

To assess iPSC-derived NK cell responses against K562 tumors cells, 1×10^5^ iPSC-derived NK cells were co-cultured with 1×10^5^ K562 cells in V-bottom 96-well plates in the presence of anti-CD107a antibodies. GolgiStop and GolgiPlug (both BD) were added after one hour to inhibit cytokine secretion. After five more hours of co-culture, cell suspensions were stained for flow cytometric analysis including fixation, permeabilization, and intracellular staining for cytokines.

To interrogate the response to innate cytokines, 1×10^5^ iPSC-derived NK cells were exposed to 10 ng/mL IL-12 (Miltenyi), 10 ng/mL IL-15, and 10 ng/mL IL-18 (both RnD) for 24 hours with GolgiStop and GolgiPlug (both BD) present during the last 5 hours, followed by staining and flow cytometric analysis.

iPSC-derived NK cells were identified as single viable CD14^−^ CD3^−^ CD56^+^ events.

### Co-culture of NK cells with allogeneic iPSC-derived NK cells

NK cell responses against allogeneic iPSC-derived NK cells were evaluated using purified NK cells from donors with specific KIR-ligand status as effectors and *B2M^−/−^ CIITA^−/−^* or *B2M^−/−^CIITA^−/−^CD54^−/−^CD58^−/−^*iPSC-derived NK cells as targets. For cytotoxicity assays, target iPSC-derived target NK cells were labelled with 2 uM CellTrace Violet (Thermo Fisher) and co-cultured with effector NK cells at varying effector:target ratios for 6 hours. Target iPSC-derived NK cells were identified as single CellTrace Violet^+^ CD3^−^ CD45^+^ CD56^+^ cells. Caspase activity, viability, and specific cytotoxicity were determined as described above.

For degranulation assays, 1×10^5^ CellTrace Violet-labelled target iPSC-derived NK cells were co-cultured with 1×10^5^ effector NK cells in the presence of anti-CD107a antibody for 6 hours. GolgiStop and GolgiPlug (both BD) were added after one hour and after five more hours of co-culture, cell suspensions were stained for flow cytometric analysis. Effector NK cells were identified as single viable CellTrace Violet^−^ CD3^−^ CD56^dim^ lymphocytes and further stratified into sub-populations as indicated.

### Co-culture of PBMC with allogeneic iPSC-derived NK cells

1×10^6^ *B2M^−/−^CIITA^−/−^*and *B2M^−/−^CIITA^−/−^CD54^−/−^CD58^−/−^*iPSC-derived NK cells were either cultured alone or co-cultured with 1×10^6^ PBMC from HLA-A2^+^ donors in complete media. CountBright Absolute Counting Beads (ThermoFisher) were used to enumerate cells every other day. iPSC-derived NK cells were identified as viable CD56^+^ HLA-A2^−^ cells and their counts were normalized to those cultured without PBMC.

### *In vivo* persistence assay of iPSC-derived NK cells in presence of activated NK cells

Purified NK cells from HLA-A2^+^ healthy donors were used as recipient compartment for these assays. To test *in vivo* persistence, 1×10^7^ unmodified, *B2M^−/−^CIITA^−/−^*, or *B2M^−/−^CIITA^−/−^CD54^−/−^CD58^−/−^*iPSC-derived NK cells were infused into female hIl15-NSG mice (Jackson) either alone or together with 5×10^6^ primary NK cells on day 0 by tail vein injection. On day 21, blood was collected and CountBright Absolute Counting Beads (ThermoFisher) were used to enumerate cells. iPSC-derived NK cells were identified as viable mouse CD45^−^ human CD45^+^ CD56^+^ HLA-A2^−^ cells and their counts were normalized to the mean of the single infusion group without co-transfer of NK cells.

**Supp. Table 1.** Antibodies.

| **Reagent** | **Source** | **Identifier** |
| --- | --- | --- |
| Anti-human α-Tubulin-AlexaFluor488 (DM1A) | Sigma Aldrich | AB_441973 |
| Anti-human CD107a-BUV395 (H4A3) | BD | AB_2739073 |
| Anti-human CD14-V500 (MΦP9) | BD | AB_2737727 |
| Anti-human CD16-BV785 (3G8) | BioLegend | AB_2563803 |
| Anti-human CD16-PerCP-Cy5.5 (3G8) | BD | AB_1727434 |
| Anti-human CD19-BV570 (HIB19) | BioLegend | AB_2563606 |
| Anti-human CD3-BV786 (UCHT1) | BD | AB_2739260 |
| Anti-human CD3-PE-Cy5 (UCHT1) | Beckman Coulter | AB_2827419 |
| Anti-human CD45-BUV395 (HI30) | BD | AB_2869519 |
| Anti-human CD54-AlexaFluor 647 (HA58) | BioLegend | AB_2715942 |
| Anti-human CD54-BV421 (HA58) | BioLegend | AB_2832690 |
| Anti-human CD54-PE (HA58) | BioLegend | AB_10897647 |
| Anti-human CD56-BUV737 (NCAM16.2) | BD | AB_2744432 |
| Anti-human CD56-BV421 (NCAM16.2) | BD | AB_2732054 |
| Anti-human CD56-BV786 (NCAM16.2) | BD | AB_2738569 |
| Anti-human CD56-ECD (N901) | Beckman Coulter | AB_2750853 |
| Anti-human CD57-PacificBlue (HCD57) | BioLegend | AB_2063198 |
| Anti-human CD57-PacificBlue (HNK-1) | BioLegend | AB_2562459 |
| Anti-human CD58-PE (TS2/9) | BioLegend | AB_1186063 |
| Anti-human CD58-PE-Cy7 (TS2/9) | BioLegend | AB_2650884 |
| Anti-human HLA class I-Alexa Fluor488 (DX17) | BD | AB_1645353 |
| Anti-human HLA class I-APC-Cy7 (W6/32) | BioLegend | AB_10709590 |
| Anti-human HLA class I-FITC (W6/32) | BioLegend | AB_314873 |
| Anti-human HLA class I-PE (DX17) | BD | AB_1645510 |
| Anti-human HLA class II-PE (Tü39) | BioLegend | AB_2750318 |
| Anti-human HLA-A2-APC (BB7.2) | BD | AB_10646036 |
| Anti-human HLA-A2-BV711 (BB7.2) | BD | AB_2740467 |
| Anti-human HLA-A2-PE (REA517) | Miltenyi | AB_2727568 |
| Anti-human HLA-E-PE (3D12) | BioLegend | AB_1659249 |
| Anti-human HLA-E-PE-Cy7 (3D12) | BioLegend | AB_2565263 |
| Anti-human CD16-PE-Dazzle594 (3G8) | BioLegend | AB_2563639 |
| Anti-human CD271-PE (ME20.4) | BioLegend | AB_2152647 |
| Anti-human CD3-BV421 (OKT3) | BioLegend | AB_2565849 |
| Anti-human CD4-BUV395 (SK3) | BD | AB_2738273 |
| Anti-human CD45-BV421 (2D1) | BioLegend | AB_2687375 |
| Anti-human CD56-BV785 (5.1H11) | BioLegend | AB_2566059 |
| Anti-human NKG2A-APC (REA110) | Miltenyi | AB_2726170 |
| Anti-human IFN-γ-AlexaFluor 700 (B27) | BD | AB_396977 |
| Anti-human IFN-γ-BV786 (4S.B3) | BioLegend | AB_2563882 |
| Anti-human IgG-Fc-PE (Goat polyclonal) | Invitrogen | AB_465926 |
| Anti-human KIR2DL1-APC (REA284) | Miltenyi | AB_2752101 |
| Anti-human KIR2DL1/S1-PE-Cy7 (EB6) | Beckman Coulter | AB_2801261 |
| Anti-human KIR2DL2/3/S2-PE-Cy5.5 (GL183) | Beckman Coulter | AB_2857331 |
| Anti-human KIR2DL3-Biotin (REA147) | Miltenyi | AB_2655344 |
| Anti-human KIR3DL1-Alexa700 (DX9) | BioLegend | AB_2130824 |
| Anti-human KIR3DL1-APC-Fire750 (DX9) | BioLegend | AB_2687394 |
| Anti-human NKG2A-Vio Bright FITC (REA110) | Miltenyi | AB_2726173 |
| Anti-human NKG2C-PE (REA205) | Miltenyi | AB_2751835 |
| Anti-human SSEA-4-FITC (MC813-70) | BD | AB_1645491 |
| Anti-human TRA-1-81-AlexaFluor 647 (TRA-1-81) | BD | AB_10550550 |
| Anti-human TNF-BV650 (Mab11) | BioLegend | AB_2562741 |
| Anti-mouse CD45-BV650 (30-F11) | BD | AB_2738189 |
| Isotype-FITC (MOPC-21) | BioLegend | AB_2884007 |
| Isotype-PE (MOPC-21) | BioLegend | AB_326435 |

**Supp. Table 2.**
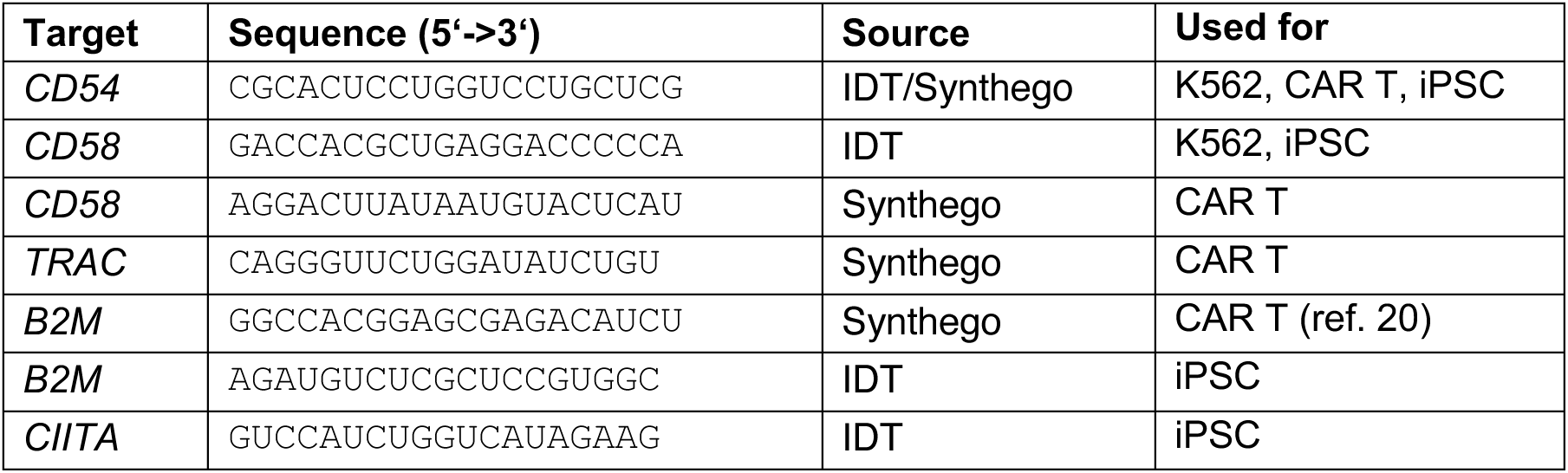
gRNA.

