## Supplemental Figures 1-4 for "Genetic ablation of adhesion ligands averts rejection of allogeneic immune cells"

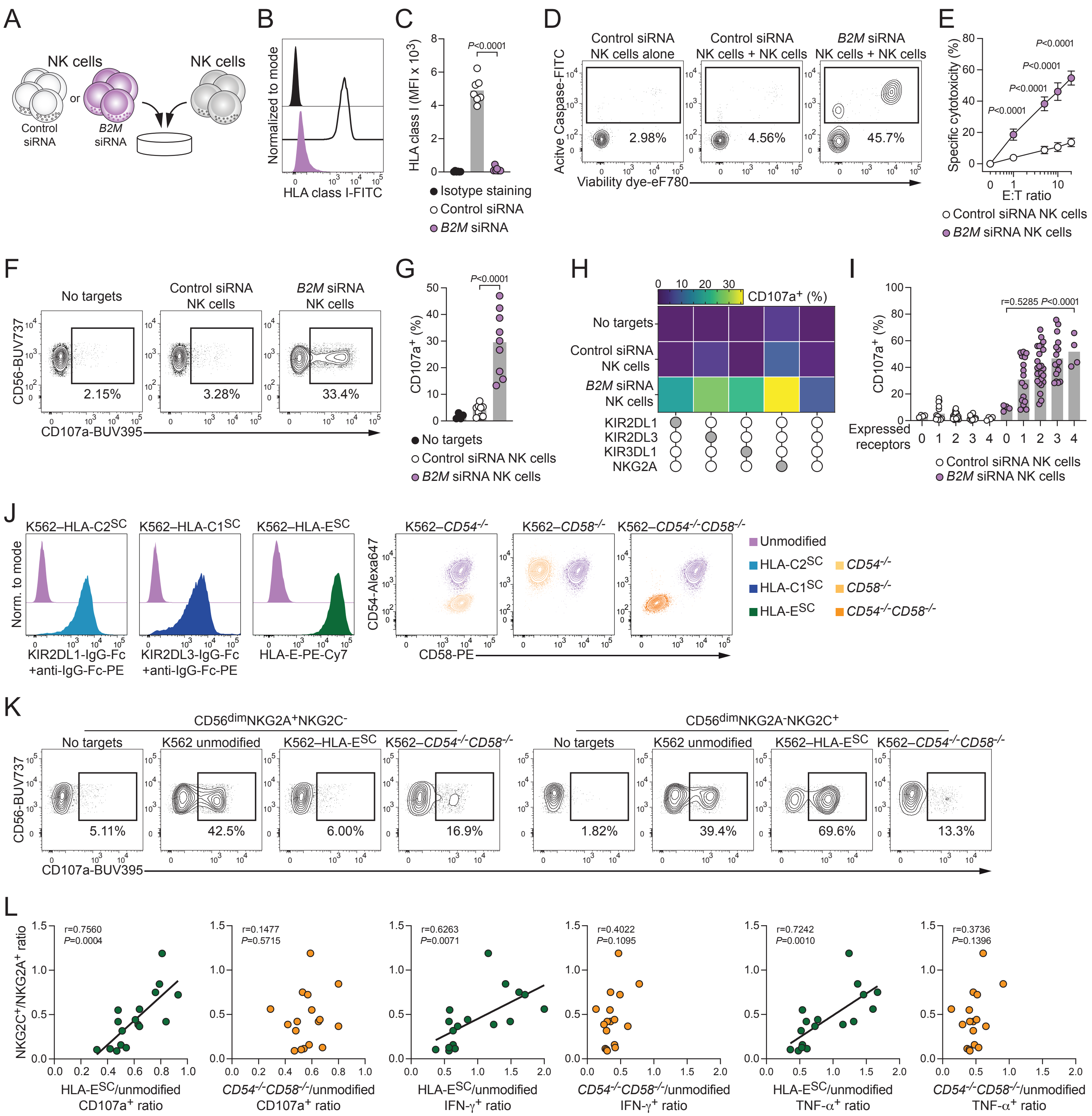


Supplemental Figure 1. Hammer *et al.*

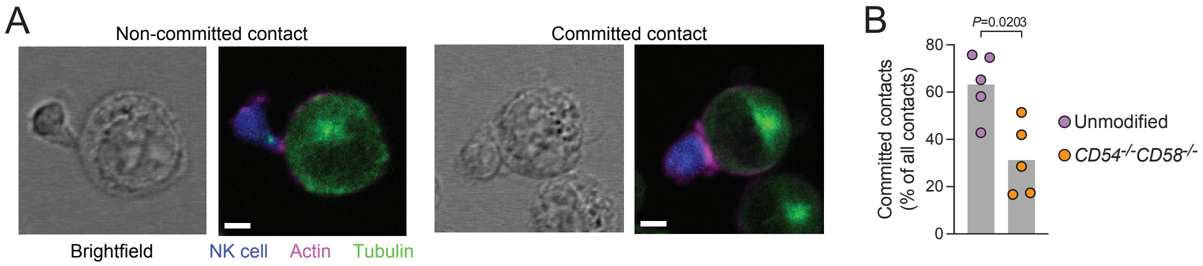


Supplemental Figure 2. Hammer *et al*.

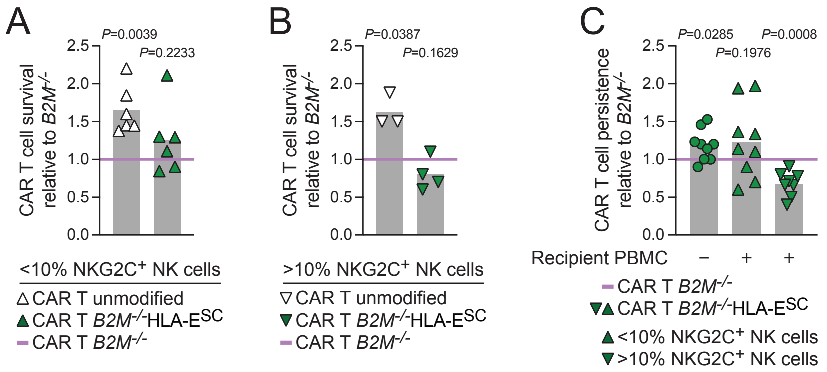


### Supplemental Figure 3. Hammer *et al*.

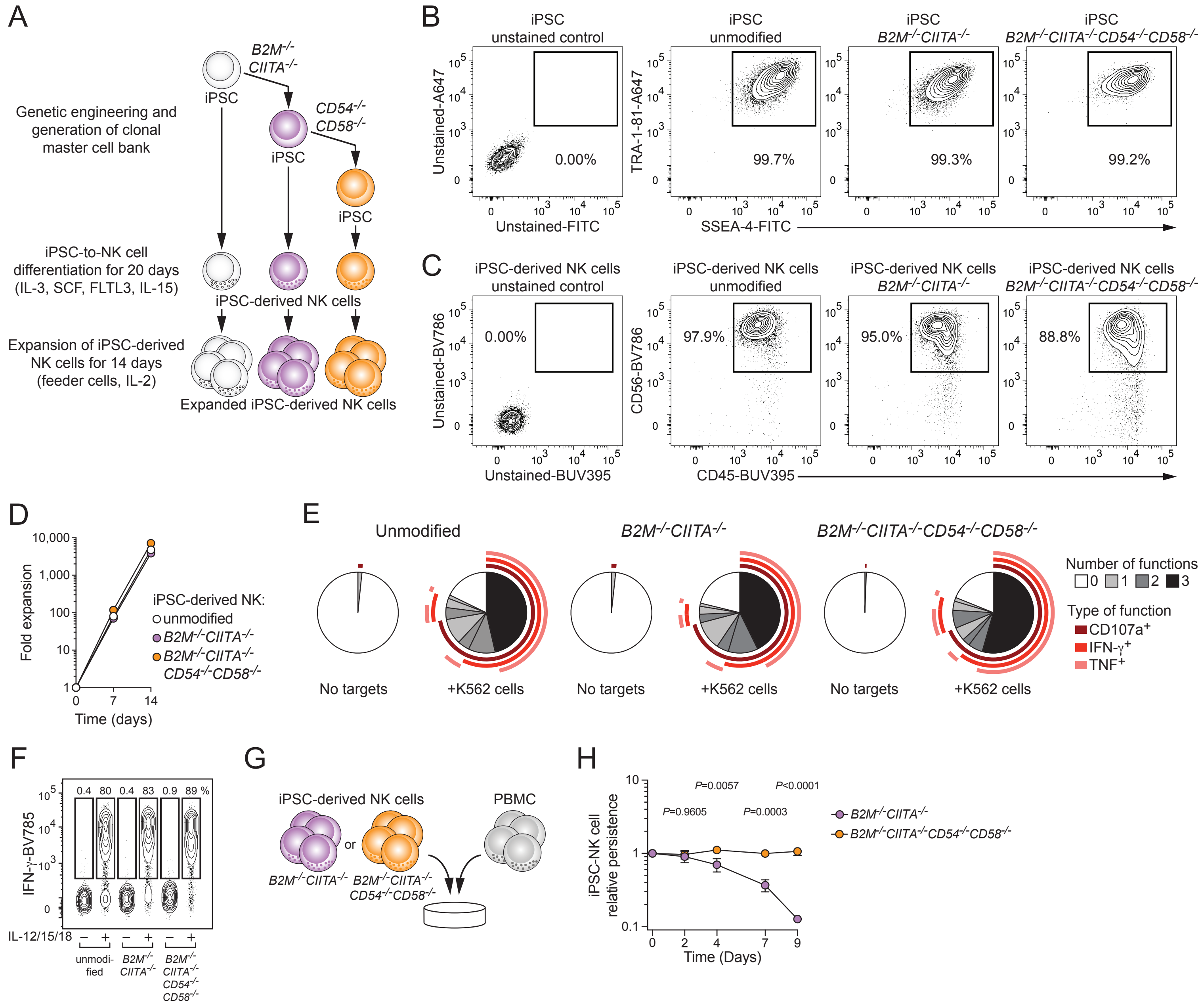


Supplemental Figure 4. Hammer *et al.*
